# Differential age effects on functional network development link to contextual memory disruption in APOE4 versus APOE3 homozygous mice

**DOI:** 10.1101/2025.05.22.655620

**Authors:** Zachary D. Simon, Karen N. McFarland, Paramita Chakrabarty, Marcelo Febo

**Affiliations:** Department of Psychiatry, University of Florida, Gainesville, FL, USA; Department of Neuroscience, University of Florida, Gainesville, FL, USA; Department of Neurology, University of Florida, Gainesville, FL, USA; Center for Translational Research in Neurodegenerative Disease, University of Florida, Gainesville, FL, USA; Advanced Magnetic Resonance Imaging and Spectroscopy Facility; Department of Pharmacology and Chemical Biology; Center for Neurodegenerative Disease, Emory University, Atlanta, GA, USA

**Keywords:** age, APOE, functional connectivity, graph theory, contextual fear conditioning

## Abstract

Apolipoprotein-ε4 (APOE4) homozygosity is the strongest genetic risk-factor for Alzheimer’s Disease (AD). The combined effects of APOE4 with age on the brain are unclear. In the present work, we tested the hypothesis that age alters contextual fear memory, functional network topology, diffusion imaging measures, and RNA expression differentially between APOE4 and APOE3 homozygous mice.

Male and female mice, 1.5-5.5 month (young) and 9-13.5 month (adult), homozygous for human APOE4 or APOE3, were scanned using an 11.1 Tesla MRI scanner. Functional magnetic resonance imaging (fMRI) and diffusion tensor imaging (DTI) were performed, and images were processed and analyzed, followed by a contextual fear conditioning (CFC) protocol to test contextual memory.

Functional imaging revealed decreases in various functional network measures and decreased network development for APOE4 adults as well as significant differences between groups in brain activity in motor, sensory, memory, and emotional processing related regions. APOE3 adult mice showed increasing network complexity with aging. DTI-based fractional anisotropy (FA) increased with age independent of genotype. Behaviorally APOE4 adult mice experienced contextual memory dysfunction relative to other groups. No sex differences were observed.

The results suggest that in APOE4 adult mice there may be a link of network connectivity changes with increases in fear behavior and a decreased ability to recognize contextual changes. Furthermore, the lack of network development in aging APOE4 mice is indicative of a loss of functional network resilience in the brains of AD-susceptible individuals.

## Introduction

Alzheimer’s disease (AD) is the most common form of dementia in people age 65 and older^1^, with ApolipoproteinE ε4 (APOE4) being the strongest genetic risk factor ^2, 3^. APOE is present in 3 unique isoforms in humans, APOE2, APOE3, and APOE4, which vary in their lipid binding capabilities and their effects on cells and other proteins in the human brain^4, 5^. Unlike the other isoforms, APOE2 and APOE3, which preferentially bind to high-density lipoproteins, APOE4 binds to large very low-density lipoproteins (VLDL) which decreases LDL receptor density, increasing LDL in the body^6–8^. Having 2 copies of *APOE*4 increases the risk of developing AD as much as 15 fold^9^. APOE4 influences several major pathways related to AD pathogenicity including increasing amyloid beta (Aβ) pathology^10, 11^, mitochondrial dysfunction^12, 13^, lipid accumulation^14, 15^, microglial dysfunction^16, 17^, and impaired neuron-astrocyte metabolic coupling^15^. In humans APOE4 has been shown to lead to hyperexcitability of frontal cortex and hippocampal neurons in addition to dysfunction and reduced signaling or a reduction in the number of GABAergic interneurons^18–21^. Understanding these and other APOE4-linked functional changes, is fundamental to finding early changes in the brain and neural/network signatures that may underlie loss of resilience and behavioral and cognitive changes in AD.

Functional connectivity (FC) in APOE4 humans has been studied largely in terms of the default mode network (DMN), considered to be a “task negative” network, most active during periods of rest^22, 23^. Lower DMN connectivity is associated with aging^24^ and older adult carriers of APOE4 that have low anti-correlated DMN connectivity have significantly lower executive function^25^. Hyperactivity in DMN and hippocampal regions in young healthy APOE4 carriers could indicate compensation of cognitive effort relative to non-carriers^26–28^. Further studies in healthy elderly APOE4 carriers found changes in DMN activity in temporal regions and the hippocampus and increases in connectivity in regions including the medial prefrontal cortex, insula, thalamus, medial temporal lobe, cingulate and striatum^29, 30^. It seems that these changes in the DMN, especially the increases in connectivity, appear in regions that eventually show significant deterioration in AD, suggesting increased cognitive burden prior to disease onset^31, 32^.Specific mechanisms of these network changes and how they affect global network function are not well understood.

Beyond the DMN there are several other networks that are consistently reported but have received less attention, especially those found in AD and regarding the role of APOE ^33, 34^. Functional neuroimaging studies of early onset AD report widespread FC declines across well establish networks^35^, including declines in graph theory based connectome measures such as network strength, efficiency and clustering in frontoinsular, parietal, temporal and occipital cortices ^36^, and reduced eigenvector centrality in the medial temporal gyrus bilaterally, right posterior hippocampal gyrus, and right postcentral gyrus^37^. However, there are disagreements across studies regarding the predominant regions, networks, and measures affected in AD^35, 38, 39^.

The use of mouse models to study APOE isoform effects on functional connectivity, genetic expression, and behavior can be used to clarify its mechanistic role in AD progression and susceptibility. Previous studies cover a range of topics from FC changes in *APOE*4tr as compared to ApoE knockout (KO) mice^40^ and dietary effects on CBF and FC in *APOE*4tr mice^41^, to effects of APOE isoform, age, and sex on cFos expression, learning behavior, and DNA damage in the hippocampus^42^. Other studies on APOE4 in mice tend to focus on neuronal and glial mechanistic changes in the brain or the effects of the removal of astrocytic APOE, often in the presence of other AD risk-factors^15, 43–47^. Two major gaps remain in these studies. First, these studies do not inform global network analysis as they lack graph theory metrics which when combined with the other methods in these papers could yield greater insight. Second, few look at an early middle-age “preclinical” timepoint. Here, using the *APOE*4tr mouse across the aging timespan from adolescence to adulthood, we found APOE4- and age-associated with white matter microstructure and functional connectome changes that relate to changes in contextual memory behavior. Further, we report gene expression changes that identify mechanistic pathways related to APOE4. We also bring to light a unique global network degradation phenotype in aging APOE4 mice. Such information could be critical to understanding the high risk conferred by *APOE*4 on developing AD.

## Methods

### Subjects and Data

Mice were housed in age- and sex-matched groups of 1-4 in a temperature- and humidity-controlled room, inside conventional air filtered cages (dimensions: 29 x 18 x 13 cm) with food and water available *ad libitum* (vivarium lights on from 07:00-19:00 hours). Mice homozygous for human APOE4 or APOE3 under the endogenous murine promoter were obtained from Duke University and generated as described in Sullivan et al. 1997 and Knouff et al. 1999^48, 49^. They were maintained as homozygotes on a C57BL6 background. APOE3 homozygous mice were used as controls for the APOE4 homozygous groups to compare the most common isoform to the most pathologically relevant isoform, and all groups had both male and female mice. Experiments were conducted in two separate age groups beginning when cohorts were between 1.5 and 5 months of age, representing a young timepoint, or between 9 and 13.5 months of age, representing a fully adult timepoint that would best represent a preclinical period in AD. The fMRI and DTI imaging group sizes were composed of 12 young APOE3, 7 adult APOE3, 12 young APOE4, and 13 adult APOE4 mice. Behavioral analysis included the same groups, minus 1 APOE4 adult that died prior to fear conditioning. RNA data of APOE3 and APOE4 targeted replacement mice was obtained from the GEO database (GSE226093) and included 9-12mo APOE3 and APOE4 homozygous mice (n=3/group). All procedures received prior approval from the University of Florida Institutional Animal Care and Use Committee and followed all applicable NIH guidelines.

### Functional Magnetic Resonance and Diffusion Tensor Imaging Instrumentation

Images were collected on a magnetic resonance spectrometer tuned to 470.7 MHz proton resonant frequency (Magnex Scientific 11.1 Tesla/40 cm horizontal magnet). The MR system was equipped with Resonance Research Inc. spatial encoding gradients (RRI BFG-240/120-S6, maximum gradient strength of 1000 mT/m at 325 Amps and a 200 µs risetime) and controlled by a Bruker AV3 HD console running Paravision 6.0.1 (Billerica, MA). An in house built 2 cm x 2.5 cm quadrature transceive surface radiofrequency (RF) coil was used for B1 signal generation and detection (RF engineering lab, Advanced Magnetic Resonance Imaging and Spectroscopy Facility, Gainesville, FL).

### Mouse imaging setup

Mice were scanned sedated under a continuous paranasal flow of 0.5 % isoflurane gas (delivered at 0.5L/min mixed with medical grade air containing 70% nitrogen and 30% oxygen, Airgas, Inc.) which was increased following functional scanning to 2.0% at 2.0L/min for the diffusion scan. Prior to setup, mice were anesthetized under 2% isoflurane and administered an intraperitoneal (i.p.) injection of 0.1mg/kg dexmedetomidine (at a 1ml/kg volume). Isoflurane was reduced to 0.5% throughout the functional imaging and increased to 2% for diffusion imaging. Functional MRI scans were collected at least 40 minutes after the i.p. injection. Respiratory rates were monitored continuously, and mice were kept warm throughout the imaging session using a warm water recirculation system (SA Instruments, Inc., New York).

### MRI acquisition protocol

We acquired one T2 weighted anatomical, an fMRI scan, and small and large b value sets of diffusion weighted scans. The T2-weighted Rapid Acquisition with Relaxation Enhancement (RARE) sequence was acquired with the following parameters: effective echo time = 41.4 ms, repetition time (TR) = 4 seconds, RARE factor = 16, number of averages = 12, field of view (FOV) of 15 mm x 15 mm and 0.8 mm thick slice, and a data matrix of 256 x 256 (0.06 µm^2^ in plane) and 12 interleaved ascending coronal (axial) slices covering the entire brain from the rostral-most extent of the anterior prefrontal cortical surface, caudally towards the upper brainstem and cerebellum (acquisition time 9 minutes 36 seconds). Functional images were collected using a single-shot spin echo planar imaging sequence with the following parameters: TE = 16 ms, TR = 1.5 seconds, 600 repetitions, FOV = 15 x 15 mm and 0.9 mm thick slice, and a data matrix of 64 x 48 (0.23 x 0.31 µm in plane) with 14 interleaved ascending coronal slices in the same position as the anatomical scan. Diffusion-weighted images were collected using a four shot 2-shell SE-EPI sequence with the following parameters: TE = 19 msec, TR = 4 seconds, average number = 2, gradient duration = 3 msec, diffusion time spacing = 8 msec, FOV = 15.36 mm x 11.52 mm, slice thickness = 0.7 mm, matrix = 128 × 96, with 17 interleaved ascending coronal slices in the same position as the anatomical scan. For DTI analysis, 71 images were collected, a B0 baseline image and 70 images with gradient directions for b values of 600 and 1800 s/mm^2^. Respiratory rates, isoflurane and dexmedetomidine delivery, body temperature, lighting, and room conditions were kept constant across subjects.

### Image pre-processing

The image processing workflow followed that of previously published research^50–53^. Resting state fMRI processing was carried out using Analysis of Functional NeuroImages (AFNI) ^54^, FMRIB Software Library (FSL) ^55^, and Advanced Normalization Tools (ANTs) ^56^. Binary masks for anatomical and functional scans were created using ITKSNAP ^57^. The brain binary masks were used for brain extraction prior to registration steps, to ensure isolation of brain voxels from images for analysis. Times series spikes were removed (3dDespike, AFNI), image repetition frames aligned to the first time series volume (3dvolreg, AFNI), and detrended (high pass filter <0.009 Hz using AFNI 3dTproject). Independent component analysis (ICA) using Multivariate Exploratory Optimized Decomposition into Independent Components (FSL MELODIC version 3.0) was used to assess structured ‘noise’ or artifact components in each subject scan, in their native space. Most, if not all ICA components in this first stage contained artefact signal voxels along brain edges, in ventricles, and large vessel regions. These components were suppressed using a soft (‘non-aggressive’) regression approach, as implemented in FMRIB Software Library (FSL 6.0.5) using fsl_regfilt ^55^. A low-pass filter (>0.12Hz) and spatial smoothing (0.4mm FWHM) were next applied to the fMRI scans prior to registration steps. Post-regression ICA was carried out to verify removal of artefact components and preliminary assessment of putative ICA networks in individual scans.

### Atlas registration and resting state signal extraction

Anatomical scans were cropped and bias field corrected (N4BiasFieldCorrection, ANTs). Functional scans were cropped, and a temporal mean image was registered to the anatomical. Preprocessed anatomical and fMRI scans were aligned to the Allen Mouse Brain Atlas,^58^ a parcellated mouse common coordinate framework (version 3, or CCFv3) template ^58^. Bilateral region of interest (ROI)-based nodes (148 total) were created with the guidance of the annotated CCFv3 parcellation and using tools in ITKSNAP and FSL, similar to our previous work in rats ^59, 60^. Large brain regions, such as hippocampus, motor, somatosensory, and visual cortices, were assigned multiple nodal masks. Thus, these are distinguished based on atlas coordinates and a numerical identifier at the end of the regional code (e.g., hippocampus contains nodes HPC1-HPC5). Anatomical images were linearly registered to the mouse template using FSL linear registration tool (FLIRT), using a correlation ratio search cost, full 180-degree search terms, 12 degrees of freedom and trilinear interpolation. The linear registration output was then nonlinearly warped to template space using ANTs (antsIntroduction.sh script). Anatomical-to-atlas linear and nonlinear transformation matrices were applied to fMRI scans at a later stage. Spontaneous signals, each with 600 data points, were extracted from the 148 ROIs and used in cross-correlations and calculations of Pearson r coefficients for every pairwise combination of ROIs (using *corrcoef* function in MATLAB). The resulting number of pairwise correlations was 10,878 per subject (after removing 148 self-correlations). Correlation coefficients were Fisher’s transformed to ensure normality prior to statistical analyses. The center voxel coordinates for the ICA-based nodes normalized to the CCFv3 were used in 3D network visualizations using BrainNet viewer ^61^.

### Network analysis

Weighted undirected matrices were analyzed using Brain Connectivity Toolbox^62^ in MATLAB (Mathworks, Natick, MA). Global graph metrics were calculated for edge density thresholds ranging from 2-40%. Global network measures for this density range were converted to area under the curve (AUC) values prior to statistical assessments. Node-specific network measures were converted to AUC values per node. We assessed several graph measures of network integration and communication efficiency APOE4/4 and APOE3/3 homozygous mice. These included node strength (sum of edge weights/node) and degree (number of edges/node) and global measures, such as transitivity (related to clustering coefficient(CC); number of triad groups normalized by all possible triad nodes in a network), and characteristic path length (CPL; the average edges or edge weights between node pairs). For local node efficiency, a length matrix was used to calculate inverse values in vicinity of a node, with added weights used to emphasize the highest efficiency node paths ^62, 63^. To corroborate results relative to random networks, all network measures were calculated on original and randomized versions of the functional connectivity matrices. Positive and negative edges were randomized by ∼5 binary swaps with edge weights re-sorted at each step. A probabilistic approach for community detection was used to calculate a modularity statistic (Q), which indexes the rate of intra-group connections versus connections due to chance ^64^. The procedure starts with a random grouping of nodes and iteratively moving nodes into groups which maximize the value of Q. The final number of modules and node assignments to each group (e.g., community affiliation assignments) was taken as the median of 100 iterations of the modularity maximization procedure ^59^. We also analyzed the tendency of assortative vs dissortative mixing of nodes ^65^1Z. The assortativity index is a correlation coefficient comparing node strength values between pairs of edge-connected nodes. Positive r values indicate connectivity between pairs of nodes with similar strengths (e.g., high strength nodes pairs with high and low with low), while negative r values indicate cross-pairings between low and high strength nodes. Functional connectivity networks were visualized in BrainNet viewer ^61^.

### Diffusion Tensor Imaging

Diffusion-weighted images were converted to NifTI and then small and large shells were combined into one image using fslmerge^55^. Images with severe noise (exhibiting high interference, random fluctuations in the signal, abnormal motion distortions) were excluded from the study. The remaining images were processed using a combination of FSL and Mrtrix software^66^. Denoising (enhancing signal-to-noise ratio), Gibbs ringing (suppressing artifactual oscillations), eddy correction (to correct for eddy current-induced image stretching, shearing, and translation), phase encoding (correcting for motion distortions), B1 field inhomogeneity (mitigating variations in the RF field) were applied to the data. These steps ensure higher accuracy and uniform signals across the brain before analysis.

### Contextual Fear conditioning

Fear conditioning was performed two weeks before fMRI studies began. The fear conditioning system and methods used were previously described^51^. The fear conditioning cages were housed in sound attenuation chambers. The chambers had computer-controlled lighting, white noise generator, a fan, and control units. These included audio speaker controls for tone generation, shock controllers for the internal cage floors, and infrared and white lighting controllers. The internal operant cage was made of translucent Plexiglas with a steel frame and steel shock grid floor connected to a computer controlled current stimulator (Med-Associates, Inc. St. Albans, VT). Activity within the fear conditioning cages was captured by a monochrome video camera. All cage accessories were controlled by PC running Video Freeze Software^TM^. Camera lens brightness, gain, and shutter settings were verified and adjusted before collecting data and were kept constant across all mice in a given chamber. All animals were tested on all days in the same box to avoid any environmental changes or effects due to differences in test cage equipment. For each session, the motion-sensing software was calibrated to 0 with the cage empty and NIR lights on. After calibration the mouse was placed in the cage for recording. The estimated motion index was set to the same threshold level (10 a.u.) across all mice prior to data exporting or analyses. Videos were acquired at 30 frames per second (fps) with a minimum freeze duration of 30 frames. For training, on the first day a 200 second baseline was followed by four presentations of a moderate level current to the entire grid floor for 1 second (0.9mA) (inescapable shock). Each of the four shock presentations were separated by 200 seconds and the entire session lasted 17 minutes. On days 2 and 3 (recall tests at 24 and 48 h of training day 1, respectively), the same protocol was carried out in the absence of the electric shock. On day 3 (modified context recall test), the cage environment was modified by placing a smooth plastic white floor and black nearly opaque diagonal walls, which covered the grid floor, light, and speaker. The cameras’ gain, brightness and contrast were adjusted to settings that could visualize the mice in this new setting and kept constant across subjects in the new chamber setting. For day one analysis, from each of the time epochs corresponding to the 4 shocks, a component comprising a 60 second pre-shock interval and a 60 second post-shock interval was used to quantify percent component freezing. On following days, the entire 17 minutes was broken into 1 minute components from beginning to end to quantify minute-by-minute changes in component freezing. Motion-index was used as a measure of locomotor activity.

### Normalized gene counts and analysis of differentially expressed genes (DEGs)

FASTQ files were downloaded from GSE226093 and aligned to the mouse genome (GRCm39) with GRCm39.107 annotation using STAR (Dobin 2013 PMID 23104886) to generate BAM files. Rsamtools and GenomicAlignments R packages ^67, 68^ were used to generate gene counts from the BAM files and were filtered for gene counts to remove genes with low counts (<10 genes). Subsequent analyses used normalized gene counts expressed as fragments per kilobase of transcript per million mapped reads (FPKM values). Analyses of DEGs was carried out using DESeq2 using default setting and a general linear model fit per gene ^69^. Wald tests were used to compare APOE4 vs APOE3 groups and multiple comparison adjustments of resulting p-values was done with the Benjamini-Hochberg false discovery rate (FDR) correction method. DEGs were defined as an absolute log2 fold change > 1 and an adjusted p value < 0.05 (visually evaluated as volcano plots). Gene annotation symbols were used to group genes according to their known expression in mouse brain cell types ^70^. Within these cell type classifications, the geometric mean of the normalized gene counts for the genes comprising the cell types was calculated and analyzed between experimental groups using Welch’s corrected t-tests. The cell type classifications included astrocytes, pan-reactive astrocytes, neurons, microglia, homeostatic microglia, disease-associated microglia, pro-inflammatory microglia, oligodendrocyte precursors, newly formed oligodendrocytes, myelinating oligodendrocytes, and endothelial cells^70–73^. Single genes were compared using normalized gene counts to determine significantly enriched and depleted genes between APOE4 and APOE3 groups using Welch’s corrected t-tests.

### Statistical analysis

Statistical analyses and data visualization were carried out using tools in MATLAB or GraphPad Prism (version 10.2). Unless otherwise stated, statistical analyses used either a two- or three-way full factorial analysis of variance (ANOVA) (genotype x age, genotype x age x trial, genotype x age x sex, critical α<0.05). Post hoc tests used Tukey’s honest significant difference (HSD) procedure. Where appropriate, FDR correction (q ≤ 0.05) was used. Further analysis of fMRI scans was carried out using probabilistic ICA ^74^ as implemented in MELODIC Version 3.15, part of FSL ^74, 75^ and as previously described for mouse fMRI scans ^51^. The resulting components were overlaid on the CCFv3 mouse atlas template and classified according to the peak z statistic anatomical location. A series of two-stage multiple linear regressions were used to back-project group ICA components to subject-specific time courses (using spatial regressions) and spatial components (using temporal regressions). The subject-specific spatial component maps were then used in statistical comparisons for each component in the FSL randomise tool. The statistical design matrix was generated using the FSL Glm tool. We used a two-way ANOVA general linear model design (age x genotype). Randomization tests were carried out with 500 permutations (corrected p-level for significance is 0.05 and statistical thresholding of maps by threshold free cluster enhancement). For DTI, we used the same Glm tool and two-way ANOVA general linear model design (age x genotype) steps to determine voxels of significantly differing fractional anisotropy, mean diffusivity, radial diffusivity, and axial diffusivity between groups.

## Results

### APOE4 adult mice exhibit increased freezing behavior during conditioning and modified context tests in a contextual fear conditioning paradigm

Three-way ANOVAs (day 1: age x genotype x trial, days 2 and 3: age x genotype x minute) of contextual fear conditioning data revealed significant differences between groups with significant effects of age (p<0.0001), genotype (p=0.0001), and trial (p<0.0001) on day 1, conditioning; age (p=0.0023) and age-genotype interaction (p=0.01) during day 2, recall; and age (p<0.0001) and age-genotype interaction (p<0.0001) on day 3, modified context. All mice develop robust fear response over the course of the conditioning time with greater freezing occurring in the following epoch surrounding each sequential shock stimulus in all groups (Figure 1a). APOE4 adult mice exhibit the greatest freezing percentage for each epoch. Day 2, young mice showed a tendency to increase freezing until about the midpoint of the testing period, after which the freezing percentage tapered off, with young APOE4 young mice consistently exhibiting lower freezing than other groups (Figure 1b). The adult mice showed unique trends in freezing with APOE3 adult mice having an earlier increase in freezing than in the young mice, followed by a larger tapering, but ending with a spike in freezing toward the end of the testing period. APOE4 adult mice showed an increase in freezing similar to the young groups, but their freezing percentage did not decrease later in the testing period to nearly the degree of other groups. On Day 3, young APOE4 mice consistently showed the least freezing when exposed to a modified context while APOE4 adult mice consistently showed the greatest freezing (Figure 1c). Furthermore, APOE4 adult mice were the only group to show consistently increasing freezing rates across the entire testing period with no large drops in freezing behavior. Trends in response curve shape in Figure 1 were reported here as the significant relationships in the three-way ANOVAs which included timing as a factor only held true on day 1 in two-way ANOVAs (age x genotype) when comparing total percent time spent freezing across the entire testing period each day. As such, with these mice, the shape of the response curve (the changes in freezing minute to minute as mice reacclimate to the space) on days 2 and 3 are important to understanding changes in contextual memory response rather than the total amount of time spent exhibiting fear behavior.

**Figure 1:**
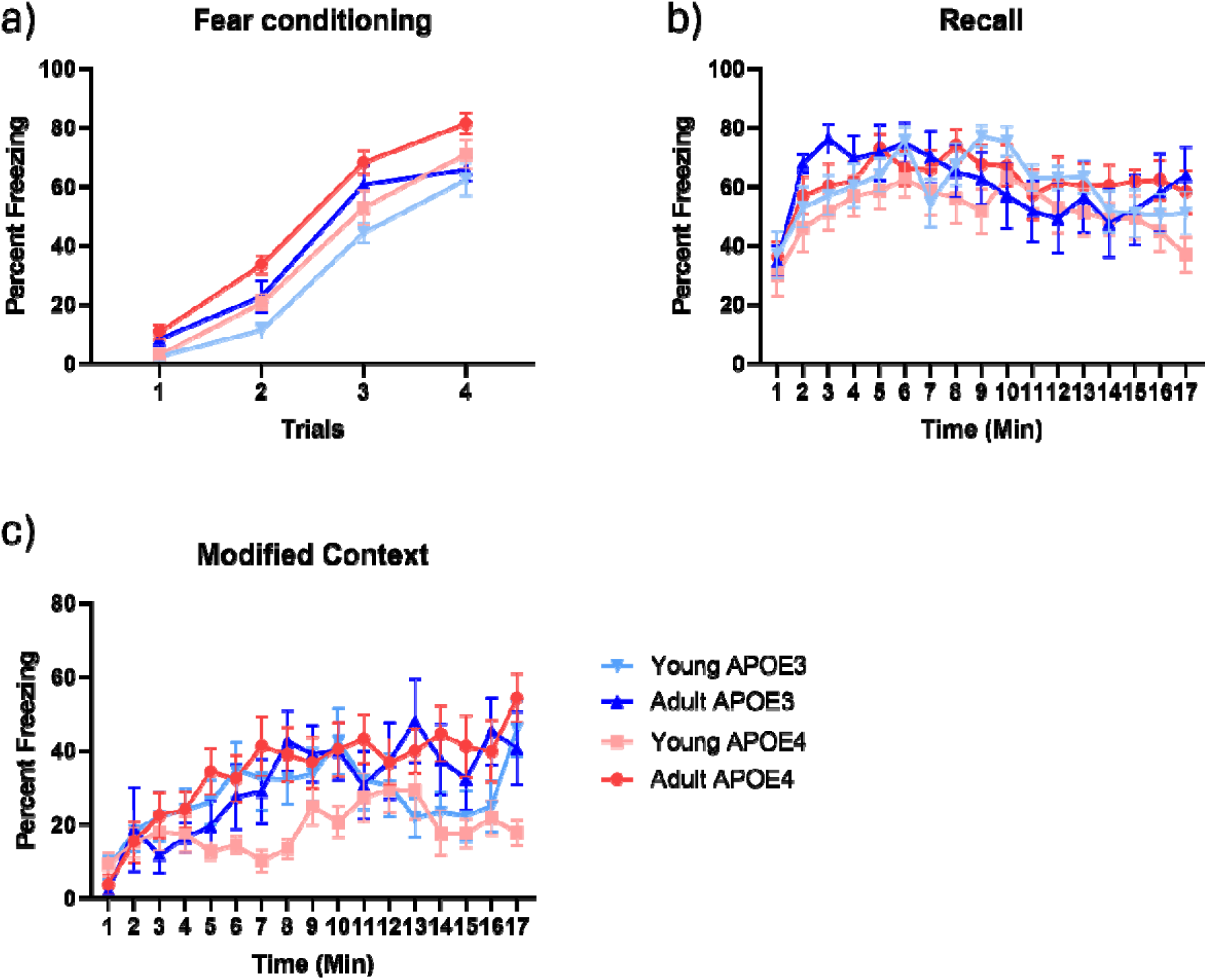
Three-way ANOVA of fear conditioning data revealed significant differences between groups during contextual conditioning tests. a) Percent freezing responses per 120 second trials (60s pre-shock, 60s post-shock) on day one of contextual conditioning with foot shock indicating robust conditioning, b) Percent freezing responses each minute over 17 minute recall session in same context without foot-shock c) Percent freezing responses each minute over 17 minute modified context recall session in modified tactile and visual context without foot-shock. Data in plots are presented as mean ± standard error

### APOE4 and age interaction decreases functional connectivity in dorsal striatal, medial visual/retrosplenial, and midbrain networks

ICA identified 20 statistically separate component networks which included previously reported networks^40, 51, 76–78^ (Figure 2). These components had peak z-statistic voxels in entorhinal, perirhinal, retrosplenial, visual, motor, and somatosensory cortical, ventral and dorsal striatal, posterior cingulate, hippocampal, amygdala, thalamic, midbrain, brainstem, and cerebellar regions. Statistical analysis using a two-way ANOVA (genotype x age) indicated a significant interaction effect of age and genotype in 4 network components with peak z-statistic voxels in dorsal striatal, medial visual/restrosplenial, and midbrain regions. APOE4 adult and APOE3 young mice had lower functional connectivity across these and other regions within these networks, than APOE4 young and APOE3 adult mice (Figure 3a). No main effects of age or genotype alone were observed in any of the 20 component networks. Figure 3b summarizes the brain areas with greater connectivity in APOE3 adult and APOE4 young mice as compared to APOE4 adult and APOE3 young mice. These included sub-regions of the motor cortex, somatosensory cortex, entorhinal, ectorhinal, perirhinal, and piriform cortices, striatum, amygdalar nuclei, reticular nuclei, insula, midbrain, brainstem, and cerebellum (p < 0.05, threshold free cluster enhanced corrected).

**Figure 2:**
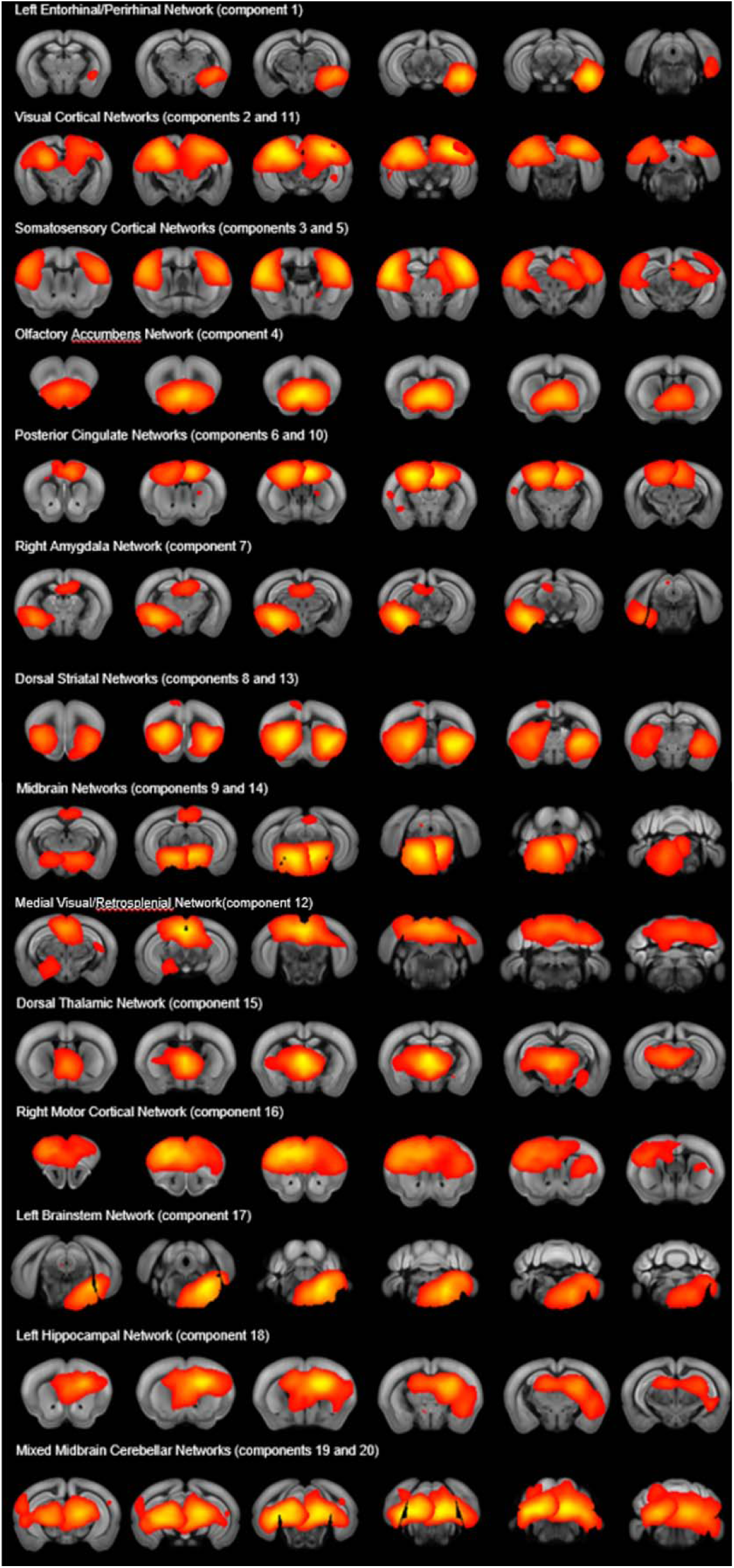
Independent component analysis of 43 mouse fMRI scans yielded 20 distinct network components. Networks were identified based on prior studies and on location of peak z statistic voxels (higher intensity/higher functional connectivity voxels highlighted in yellow and lower intensity in orange). Assigned network names are above each row of ICA maps, with component numbers in parenthesis. ICA maps are organized rostral to caudal from left to right. Z statistical maps are thresholded using threshold free cluster enhanced approach and are overlaid on to the Allen Mouse Brain Common Coordinate Reference atlas (CCFv3).

**Figure 3:**
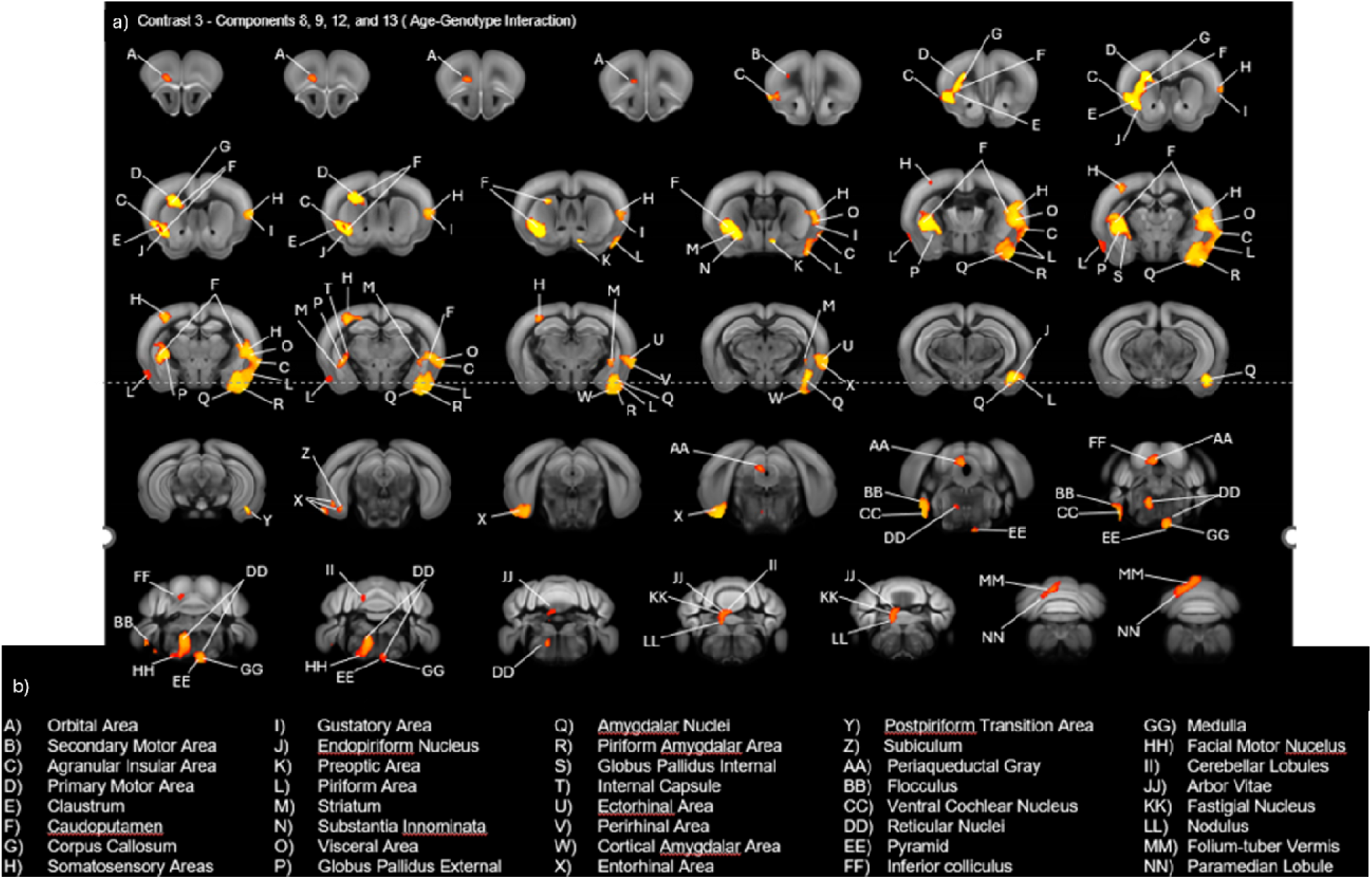
Statistical results of full factorial ANOVA assessing genotype x age interactions and main effects (p < 0.05, corrected). APOE4 adult decreased functional connectivity with a main effect of age-genotype interaction within 4 classified components in 3 networks: the dorsal striatal network, medial visual/retrosplenial network, midbrain network. This activity difference included nodes in the striatum, amygdala, and various cortical regions. Letter codes indicate the brain area according to the Allen Mouse Brain Common Coordinate Reference atlas (CCFv3). Brain area names are matched to letter codes at the bottom of the figure. Significant voxels (main effect age-genotype interaction) are highlighted in yellow-orange with yellow indicating peak z-statistic voxels for higher intensity/higher functional connectivity.

### APOE4 and age interaction is associated with lower brain-wide graph theory metrics of network connectivity and network degradation in functional connectivity network maps

Figure 4a shows graph theory metrics calculated as area under the curve for network edge-densities from 2%-40%. Global network measures of efficiency (p=0.0288), clustering coefficient (p=0.0114), transitivity (p=0.0209), assortativity (p=0.0423), and network strength (p=0.0116) varied significantly with an age-genotype interaction effect, with lower values seen in APOE4 adult mice (Figure 4a). Modularity showed a significant main effect of genotype (p=0.0414). Network maps indicating connectivity between our 148 a priori nodes, visualized in BrainNet Viewer, with edges shown above a 10% edge-density threshold, exhibited similar levels of development between APOE3 and APOE4 young mice with slightly more robust networks in APOE4 young mice. APOE3 adult mice showed greater development in their networks as compared to APOE3 young mice, indicating increased complexity and greater connectivity between more regions over the course of aging. However, APOE4 adult mice showed network degradation rather than development as compared to APOE4 young mice with reduced complexity and number of network connections over the course of aging.

**Figure 4:**
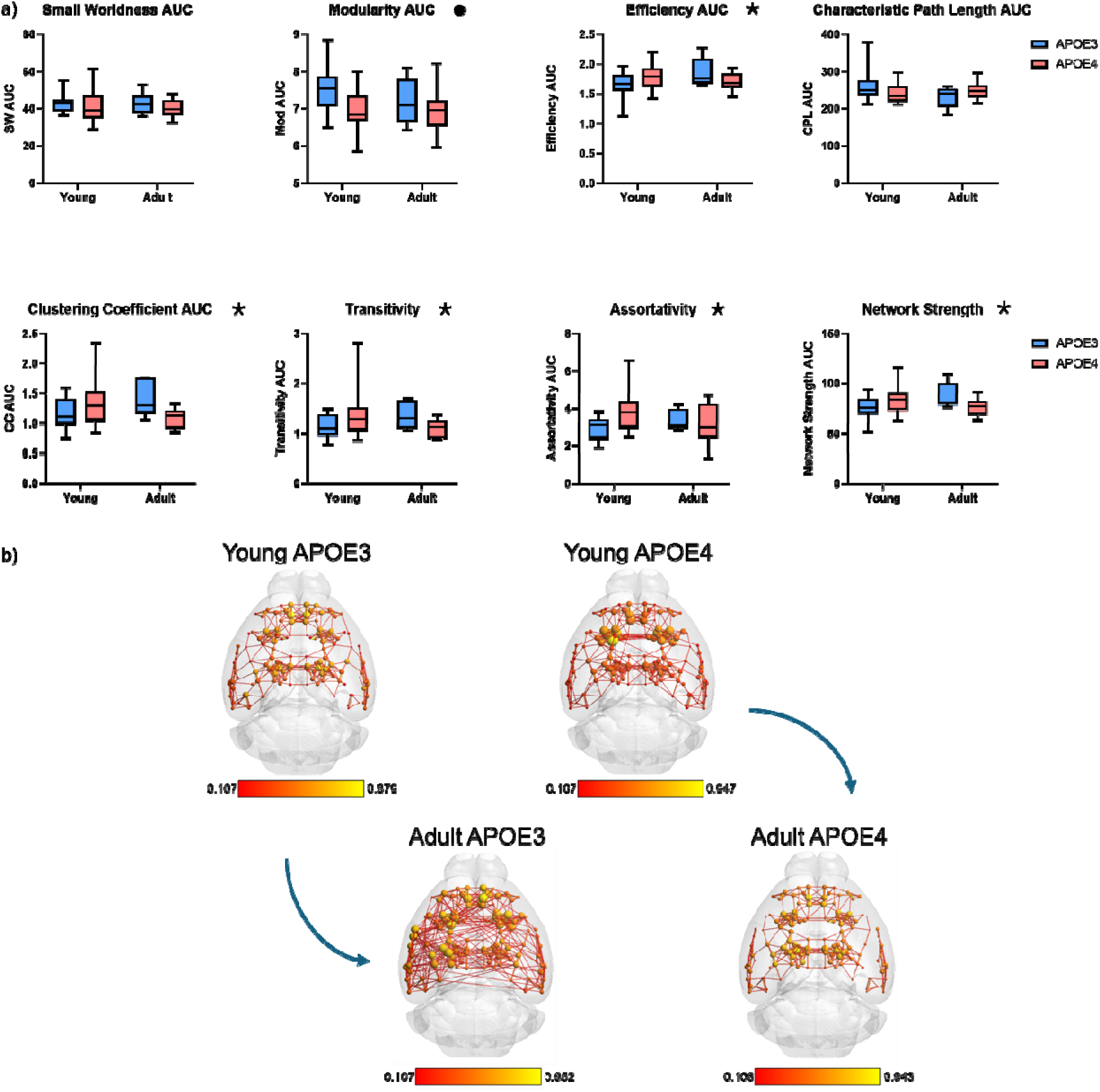
Significant differences in globally estimated graph theory metrics. a) Comparison of graph theory-based network measures (converted to area under the curve for network densities between 2-40%) between young and adult, APOE3 and APOE4 mice. Asterisks indicate metrics with significant age-genotype interaction. • indicates metric with significant main effect of genotype. b) Functional connectome maps averaged within groups and illustrate node strength and edge strength (Pearson correlations) as spherical nodes and lines connecting nodes. Node strength is reflected in sphere size and color intensity (scale bar below maps indicate Fisher z-transformed r values rescaled between 0 and 1). Edges are thresholded at 10%. Illustrates differences in development of functional networks between APOE3 and APOE4 mice in aging.

### Age increases fractional anisotropy across APOE genotypes throughout the brain in the targeted replacement APOE mouse model

Diffusion tensor imaging revealed a significant effect of age on fractional anisotropy (FA) throughout the brain (Figure 5) using a two-way ANOVA (genotype x age) (p < 0.05, corrected). However, there was no main effect of genotype or age-genotype interaction. The regions with significantly greater FA in adult mice were not localized to specific regions or structures, though many significant voxels were present in white matter. This indicates that white matter compactness and uniformity increases with aging alone regardless of APOE isoform. Significant differences with main effects of age, genotype, or interaction effects in other diffusion imaging metrics including axial diffusivity, radial diffusivity, and mean diffusivity were not observed.

**Figure 5:**
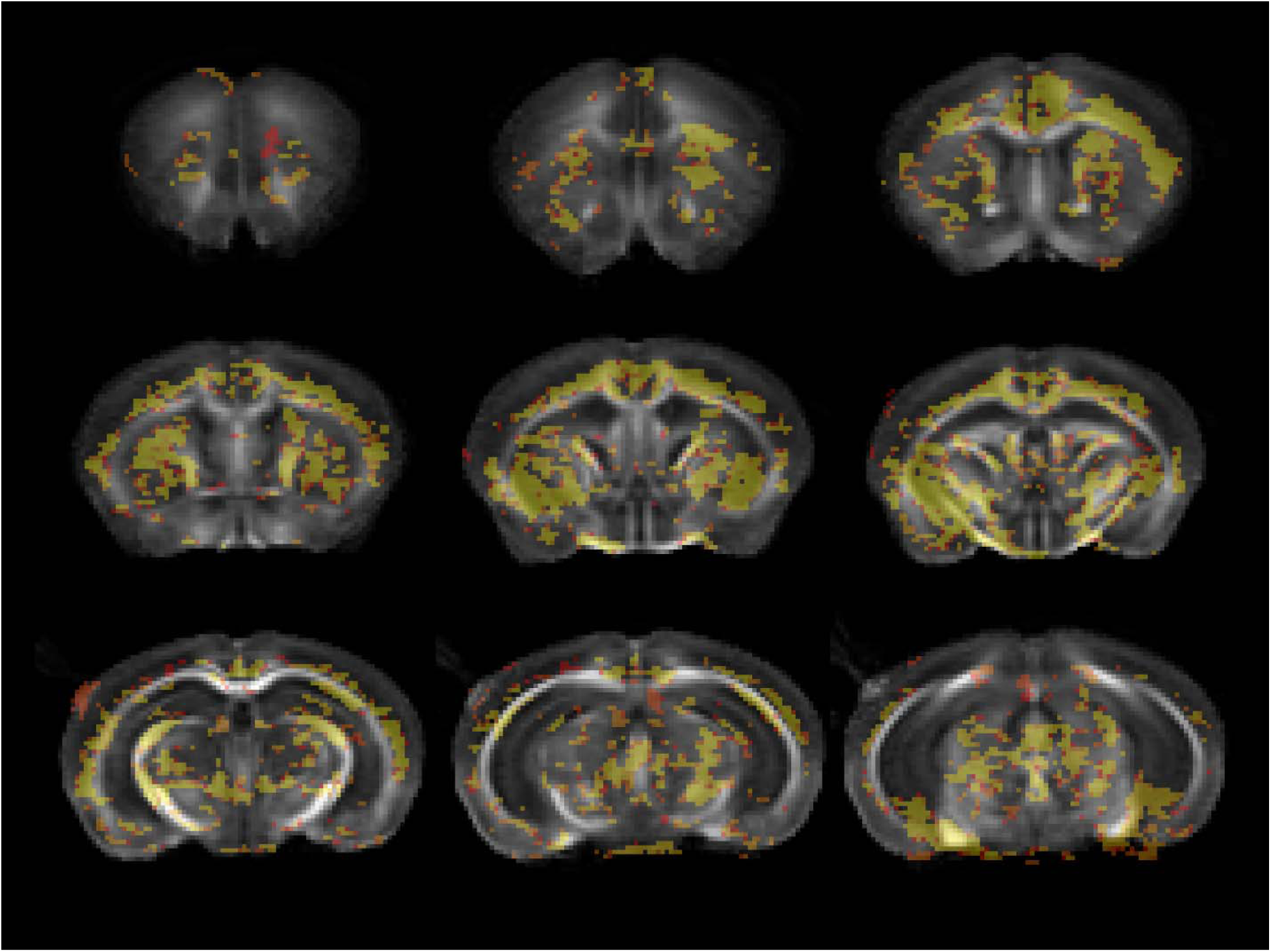
Statistical results of two-way ANOVA assessing main effect of age in fractional anisotropy (p < 0.05, corrected). FA increases with age regardless of APOE isoform genotype. Regions of significantly increased FA are highlighted in yellow-orange with yellow indicating peak z-statistic voxels for higher intensity/higher FA values. These regions are diffuse throughout the brain. Left-right, top-bottom indicates rostral-caudal sections.

### APOE4 mice exhibit significantly enriched and depleted cell type specific RNA signatures and RNA expression in genes of interest

DEG analysis revealed significant differences in cell type specific RNA signatures between APOE3 and APOE4 mice. By geometric mean of normalized cell counts for cell type specific gene groups, APOE4 mice showed significant enrichment in genes specifically related to pan-reactive astrocytes (p=0.0078, Welch’s corrected t-test) as classified by Liddelow et al. in 2017^71^ (Figure 6a), and significant depletion in endothelial cells (p=0.0185, Welch’s corrected t-test) and microglia (p=0.0491, Welch’s corrected t-test) as classified by Zhang et al. in 2014^70^, and homeostatic microglia (p=0.0231, Welch’s corrected t-test) as classified by Sala Frigerio et al. in 2019^72^ (Figure 6b-d). No significant differences were observed in other cell type specific classifications. Specifically, enriched genes Serpina3n (p=0.0044, Welch’s corrected t-test) and depleted genes Hspb1 (p=0.0226, Welch’s corrected t-test) and CD44 (p=0.0001, Welch’s corrected t-test) related to pan-reactive astrocytes^71^ (Figure 7a), were found to have significant differences in expression. For genes related to microglia^70^ (Figure 7b), Lrrc25 was significantly enriched (p=0.0016, Welch’s corrected t-test), and Pltp (p=0.0277, Welch’s corrected t-test), Cxcl16 (p=0.0481, Welch’s corrected t-test), C1qa (p=0.0384, Welch’s corrected t-test), and H2-K1(p=0.0032, Welch’s corrected t-test) were significantly depleted. Significantly enriched and depleted genes, in APOE4 as compared to APOE3 mice, not included in the above cell type groups were reported in Table 1.

**Figure 6:**
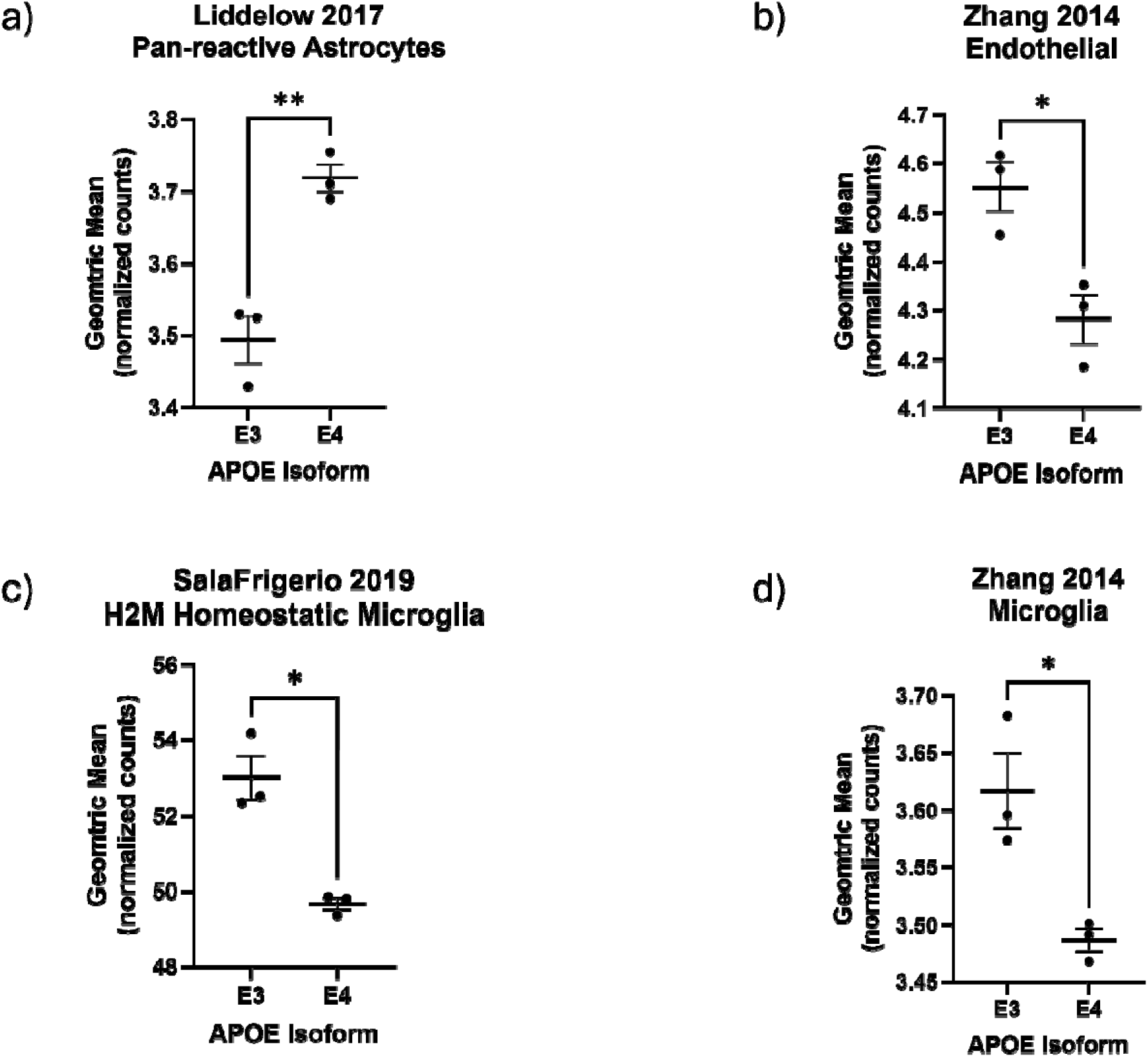
Results of RNA-sequencing. Significantly enriched (a) or depleted (b-d) cell-type specific RNA-based gene sets between APOE3 and APOE4 mice, by geometric mean of included genes. Violin plots show mean and standard error. *p<0.05, **p<0.01.

**Figure 7:**
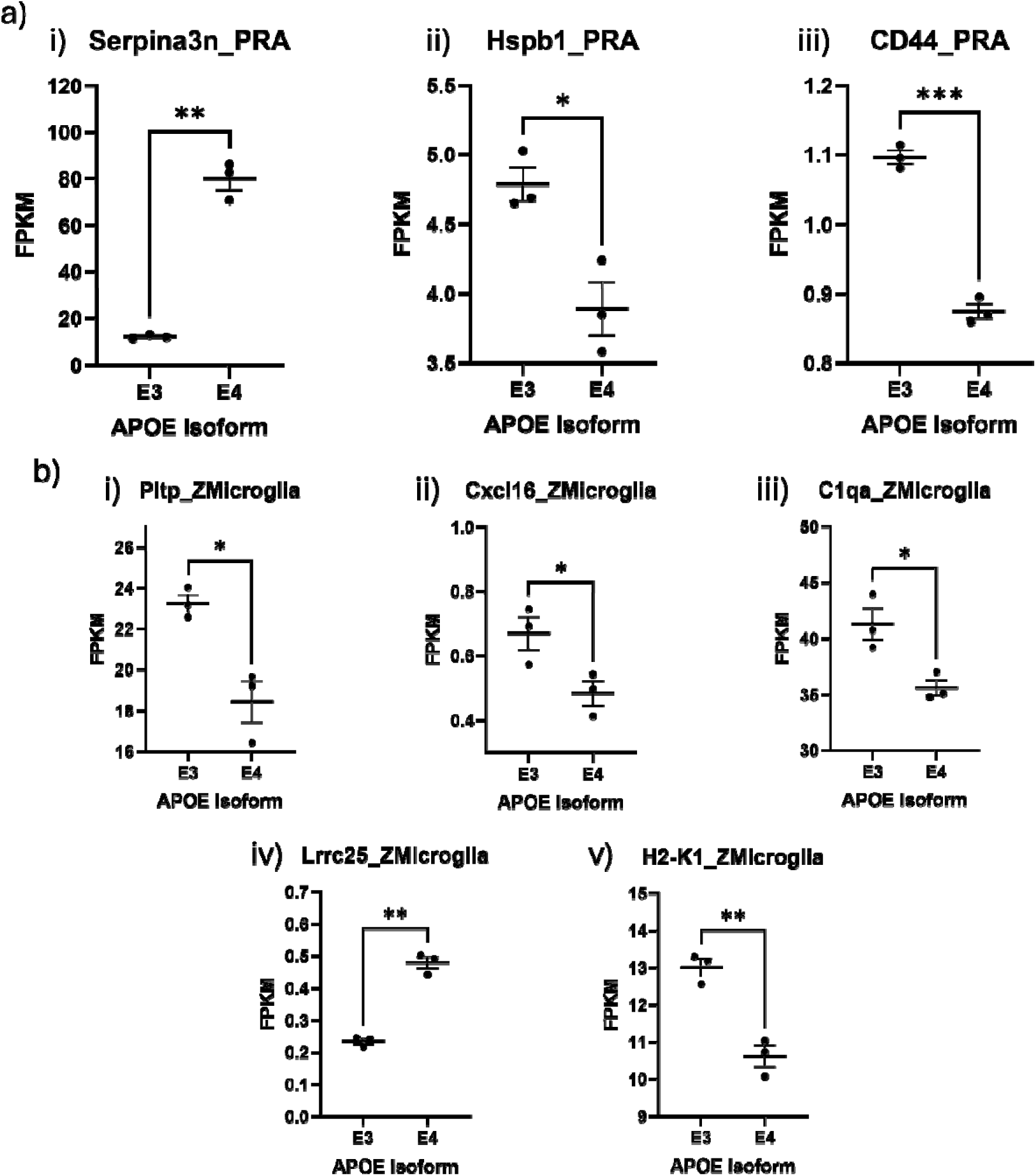
Results of bulk RNA-sequencing on APOE3 and APOE4 mice. In APOE4 as compared to APOE3 mice significantly enriched (i) or depleted (ii,iii) single genes contributing to (a) pan-reactive astrocytes (Liddelow, 2017) and enriched (iv) and depleted (i,ii,iii,v) single genes contributing to (b) microglia (Zhang, 2014) in APOE4 as compared to APOE3 mice, by fragments per kilobase million (FPKM). Violin plots show mean and standard error. *p<0.05, **p<0.01, ***p<0.001.

## Discussion

In this study we examined how APOE4 homozygosity alters brain-wide functional connectivity, white matter microstructure, contextual fear memory, and forebrain gene expression in young and adult APOE3 and APOE4 targeted replacement mice. Our data show brain connectomic changes, supported by behavioral and functional connectivity alterations that elucidate the effects of APOE4 as a risk-factor for neurodegenerative and cognitive disease. While these mice do not have AD related neuropathology (amyloid plaques or tau neurofibrillary tangles) they express human APOE4, which has been shown to be sufficient to lead to aberrant neuronal, astrocytic, and microglial activity^11, 79^. Based on evidence of cellular functional and mechanistic changes in APOE4 mice^3, 10–17^, functional changes in APOE4 humans, and the high risk of AD associated with the APOE4 isoform^3, 80^, in addition to the reversal of cognitive ability seen over the course of aging in APOE4 harboring individuals as compared to those with APOE3^81–85^, we hypothesized significant differences in brain-wide functional connectivity as a result of the age and APOE4 interaction, even between young and middle-aged mice, though previous studies from other groups tend to focus on older timepoints. As hypothesized, we found significant differences in functional connectivity with a primary effect of age-genotype interaction in established mouse ICA networks. These effects within these networks were observed in regions involved in motor and sensory processing, emotional regulation, memory, and contextual processing including amygdalar nuclei, somatosensory and motor cortex, striatum, perirhinal, entorhinal, and piriform areas. This possibly relates to the loss of synaptic density previously reported in these mice^86^, also shown in AD patients to impair functional and structural connectivity^87^. These regions may be selectively affected due to possible variations in astrocytic and neuronal expression of APOE4, or could be affected earlier in the lifespan by APOE4 than other regions associated with AD. Interestingly, to an extent, the regions with significant differences in activity in this study mirror those regions effected in earlier stages of AD by tau progression^88^. It will be important for future studies to verify the regional progression of the effects APOE4 and how these changes in brain function and how these changes relate to neuronal activity on the small circuit scale and larger network scale.

Our network connectivity maps revealed a unique previously unreported connectivity pattern resembling network degradation in APOE4 adult mice, which could underlie the risk-increasing propensity of this isoform. More robust networks tend to be more integrated and highly connected as measured by clustering and transitivity and network strength. These stronger networks are possibly related to improved cognitive ability as changes in many graph theory-based network measures accompany cognitive loss in AD^89, 90^. Our young APOE4 mice show slightly more robust networks than their APOE3 counterparts, which is consistent with cognitive ability in human literature^81–85^. However, while APOE3 mice develop robust and complex networks as they age, APOE4 mice do not show development, but rather exhibit degradation of their network complexity and degree of connections. These findings are supported by our age-genotype significant differences in graph theory connectomic metrics. This global network-level difference in complexity and connectivity could underlie AD risk from APOE4. If a robust network is affected at the node level by an insult that leads to local functional connectivity loss, such as synaptic and neuronal damage or loss and communication deficits due to amyloid-β oligomers or plaques, or tau neurofibrillary tangles, the network should be able to compensate for this functional loss. However, in a sparse network, the same insult leading to local loss might not be easily compensated for by other connections between regions that are functionally connected. It stands to reason that the brains of APOE4 homozygotes would be more vulnerable to the effects of other risk-factors of AD, such as amyloid and tau, and that the negative changes in the brain associated with these risk-factors would be exacerbated by APOE4 homozygosity^91–94^. This could be a macroscopic reason that APOE4 is such a tremendous risk-factor for AD, though directly linking molecular mechanisms to how they affect network-wide functional connectivity and connectomic outcomes would be necessary to understand what leads to these changes beyond the general knowledge of cellular and metabolic changes due to APOE4.

Our behavioral results showed that adult APOE4 homozygous mice are more sensitive to the conditioning process and exhibit a greater fear response with main effects of genotype and age. This is likely due to an underlying change in the hippocampus-amygdala circuit that is responsible for encoding contextual information and producing a fear response in addition to linking the context to the response. It is possible that greater activity or lower inhibition in this circuit produces this effect, as it has been shown that greater amygdala activity is directly associated with increased fear response and greater hippocampal activity with contextual memory^95^. A potential mechanism of interest underlying our results are changes to GABAergic neurons in the hippocampus and amygdala are impaired which leads to overactivity of the excitatory neurons in these regions as hyperactivity of neurons has been seen in previous studies of APOE4^19, 96^. Recall of contextual fear relies on the same hippocampus-amygdala circuit as encoding fear, but different parts of the hippocampus are more involved in retrieval as compared to encoding^97^. The main effect of age indicates that over the time course adult mice show greater fear response than young mice. This could be due to greater activity in the subiculum of older mice than younger mice.

From our modified context experiment we see that age affects the ability of these mice to process a changing context, as does the age-genotype dependent interaction. This could indicate a change in the both hippocampus and prefrontal cortex responsible for recognizing and adapting to new environments based on prior experience^98^. With our younger APOE4 mice we see that we would expect somewhat normal exploratory behavior in the modified environment (intermittent amount of freezing and exploring, not dependent on time with no significant increase or decrease over time). However, with our adult groups and with the age-genotype interaction, we see a significant increase in the amount of freezing across the time-course for adult and especially adult APOE4 mice, indicating an inability to properly recognize and react to a contextually modified environment. This fear behavior generalization, or failure to discriminate contextual change, only occurs in healthy mice if exposed to the modified context soon after training, not usually after 48 hours as we are seeing here^99, 100^. This unique modified context phenotype of not just greater fear as usually seen in generalization, but an increasing level over the time spent in the modified environment points to possible aberrant frontocortical, hippocampal and amygdala activity, though this needs further exploration to understand the mechanism and significance of this finding. Furthermore, total freezing did not differ drastically between groups without the minute-by-minute effect considered. In looking at contextual fear conditioning it may be important to recognize the differences in the trajectory or shape of the fear curve across the experimental timeframe, rather than simply the total amount of freezing behavior. It is important to note the regional differences thought to underlie behavior may relate to but not exactly align with the changes we see in ICA based analysis as the fMRI is conducted at rest under light sedation. Nevertheless, some of the same regions are implicated in both.

Aging also has been shown to effect cognition and behavioral characteristics on its own in addition to increasing the production of APOE^101–103^. While not dependent upon each other during conditioning, there was a synergistic effect of aging and APOE4 on contextual fear development in this study, indicating that age and APOE genotype may work together to produce greater effects on the brain and subsequent behavior before recall. The age-genotype interaction effect on fear generalization in the modified context recall sessions implicates a dependent synergistic effect when this circuit is activated in recall as compared to encoding. These outcomes align with APOE4 and age being two of the greatest risk factors for AD. If together they induce greater changes in the brain, then what we see in AD could be partially due to this synergism and its relationship with other risk-factors. As these factors create an environment in which Aβ and tau aggregation progress quickly^104^, actively increasing amyloid plaque formation^10^ and tau-mediated neurodegeneration^47^, understanding the underlying mechanisms and finding ways to decrease these effects could be important to treating and preventing AD.

DEG analysis revealed that differences we observed in connectivity and behavior could be reliant on cell type specific changes in glial function, particularly pan-reactive astrocytes and microglia. Analysis of significant gene enrichment and depletion in APOE4 as compared to APOE3 mice pointed to key genes related to inflammation and immune activity^105–110^, lipid metabolism^107, 111, 112^, and homeostasis^106, 113^, which contributed heavily to the cell type specific differences between these groups. Furthermore, some of these genes are directly related to human AD and these genetic differences in APOE4 homozygotes could point to further mechanisms by which APOE4 induces risk to AD. In humans the molecular chaperone HSPB1 is secreted by astrocytes and has protective functions in the brain^106^, specifically related to tau and amyloid. As its mouse homolog is significantly depleted in APOE4 mice from our analysis, it is possible that this alteration increases brain related vulnerability. Furthermore, as Hspb1 plays a role in astrocytic and neuronal signaling, it could contribute to the functional connectivity changes we observed. Another gene that showed enrichment in our analysis, Lrrc25, has recently been suggested as an AD risk-factor^114^. The protein coded by this gene has been shown to be increased in AD mouse models, as has its human homolog in post-mortem human AD samples^115^. While it’s mechanism in AD is currently unknown it has been suggested that it may play a role in increasing amyloid plaques and tau neurofibrillary tangles^115^. The other genes differentially expressed by either astrocytes or microglia in our analysis, with implications in lipid metabolism and inflammation could also influence functional connectivity. Microglial cells play a phagocytic role in synaptic pruning ^116^, particularly of unused or damaged dendritic spines ^117^, which may contribute to optimization of neural circuit processing ^116, 118–120^. Astrocytes also play a crucial role in maintaining synaptic and overall neuronal function, signaling, and health^121–125^. Thus, it stands to reason that functional alterations in these glial cell populations could impart reduced synaptic integrity and neuronal signaling. Furthermore, lipids play yet another crucial role in synaptic function and lipid accumulation and dysfunction can interfere with neuronal and glial signaling and function as seen throughout APOE4 literature^12, 15, 44, 126, 127^. Overall, our DEG findings provide support for our functional connectivity findings, and provide further links between the effects of APOE4 and AD.

In conclusion, the present study adds to the literature on the functional network, contextual memory, and genetic expression changes in response APOE4 homozygosity. The behavioral assay points to GABAergic tone loss as a possible factor involved in cognitive changes seen in APOE4. The functional connectivity analysis reveals a unique global network-level developmental deficiency that could be important to understand how APOE4 confers risk and vulnerability for AD. This work indicates that APOE4 imparts changes in the brain even in early adulthood, and thus treating the issues caused by APOE4 early in life could significantly reduce risk for developing AD at later timepoints. Furthermore, these findings could be used as a basis for looking into new ways to mechanistically alter the action of APOE4 to normalize network development to possibly reduce or prevent AD risk from APOE4. It will be important to study changes at a synaptic level to further elucidate the mechanistic effects of APOE4 and how these lead to large-scale changes in functional connectivity, and to determine ways in which these alterations could be reduced. These mechanisms could be important to developing new treatments for APOE4 induced AD-risk, lowering the number of AD cases that develop long-term.

## Acknowledgements

Marcelo Febo and Zachary Simon are thankful for the support provided by the McKnight Brain Institute of the University of Florida.

## Author contributions

Zachary Simon (Data curation; Visualization; Writing - Original Draft Preparation; Writing – Review & Editing; Conceptualization; Methodology; Formal Analysis; Investigation); Karen N. McFarland (Writing - Review & Editing; Methodology; Formal Analysis); Paramita Chakrabarty (Writing - Review & Editing; Conceptualization); Marcelo Febo (Writing – Review & Editing; Conceptualization; Methodology; Formal Analysis)

## Statements and declarations

### Ethical considerations

All procedures received prior approval from the Institutional Animal Care and Use Committee of the University of Florida and follow all applicable NIH guidelines.

### Consent for publication

All authors approve of the final version of this manuscript.

